# Backscattering Mueller Matrix polarimetry on whole brain specimens shows promise for minimally invasive mapping of microstructural orientation features

**DOI:** 10.1101/2022.08.15.503997

**Authors:** Justina Bonaventura, Kellys Morara, Rhea Carlson, Courtney Comrie, Noelle Daigle, Elizabeth Hutchinson, Travis W. Sawyer

## Abstract

Understanding microscale physiology and microstructural cellular features of the brain is key to understanding the mechanisms of neurodegenerative diseases and injury, as well as the prominent changes that neurons and glia in the brain undergo in development and aging which reflect functional state. Non-invasive imaging modalities sensitive to the microscale - especially diffusion magnetic resonance imaging (dMRI) - are extremely promising for three-dimensional mapping of cellular microstructure of brain tissues and brain connectivity via tractography; however, there is a need for robust validation techniques to verify and improve the biological accuracy of fiber orientation information derived from these techniques. Recent advances in dMRI acquisition and modeling have moved toward probing of the more complex grey matter architecture, challenging current validation techniques, which are largely based on *ex vivo* staining and microscopy focusing on white matter. Polarized light imaging (PLI) has been shown to be a successful technique for high resolution, direct, microstructural imaging and has been applied to dMRI validation with clear advantages over conventional staining and microscopy techniques. Conventionally, PLI is applied to thin, sectioned samples in transmission mode, but unlike histologic staining, PLI can be extended to operate with high sensitivity in reflectance mode and even extended to 3D imaging to bridge the gap toward *in vivo* validation of dMRI measurements of orientation features in both gray and white matter of the brain. In this report we investigate the use of backscattering Mueller Matrix polarimetry to characterize the microstructural content of intact Ferret brain specimens. The experimental results show that backscattering polarimetry can probe white matter fiber coherence and fiber orientation in whole brain specimens, and show promise for probing grey matter microstructure. Ultimately, these preliminary results motivate further study to fully understand how backscattering polarimetry can best be used for validation of *in vivo* microstructural imaging of the brain.

## 1 INTRODUCTION

Developing methods to probe microscale physiology and microstructural cellular features within the brain is a rich topic of research both in the context of clinical care and neuroscientific studies. Understanding structure-function relationships in the human brain is critical both to understanding normal brain development and function, and also to elucidate the mechanisms of disorders and diseases that alter brain physiology, for example neurodegenerative diseases such as Alzheimer’s and Parkinson’s disease (Lamptey et al. (2022)).

Advances in neuroimaging technologies have enabled a wide range of research studies to explore changes in the brain structure and function during healthy aging and under pathological conditions. In particular, the last two decades have seen an explosion of studies using Magnetic Resonance Imaging (MRI), which enables *in vivo,* depth-resolved, and minimally invasive 3D visualization of structural and functional properties of the brain (Wang et al. (2021); Forouzannezhad et al. (2019)). Diffusion MRI (dMRI) techniques, such as diffusion tensor imaging, are a subset of MRI that generate contrast using the diffusion of water molecules (Novikov et al. (2019)). dMRI can be used to create exquisite maps of spatial organization and connectivity of white matter, as the cell structure of axons limits molecular movement perpendicular to the axonal fibers, giving rise to anisotropic diffusion (Le Bihan (2014)). As such, dMRI has become the modality of choice for investigating the neural architecture of the brain.

dMRI has been used extensively to study white matter changes during brain development and aging (Rathi et al. (2014); Gunning-Dixon et al. (2009); Madden et al. (2009)), in the context of pathological disorders and diseases (Nguyen et al. (2022); Zhao et al. (2022); Rosenkranz et al. (2022)), as well as from traumatic brain injury (Hutchinson et al. (2018); Minchew et al. (2022); Guerrero-Gonzalez et al. (2022); Liu et al. (2022)). However, the application of dMRI to cortical gray matter analysis has only recently begun to be explored, with studies reporting age related changes using diffusion imaging based measures (Bhagat and Beaulieu (2004); Pfefferbaum et al. (2010)). While dMRI is a powerful technique, one technological limitation is a lack of sufficient specificity to directly measure microstructural and molecular quantities, instead relying heavily on biophysical modeling or indirect inference; as a result, validation of the ‘‘biological accuracy” of dMRI estimates is essential for advancing the technology. Validation of dMRI metrics most commonly utilize histologic staining which is limited by stain specificity, tissue processing and sectioning artifacts, and a lack of quantitative metrics. While more sophisticated methods including structural tensor analysis and electron microscopy have improved these efforts, these all remain limited to the study of fixed, sectioned, *ex vivo* tissues with mm-scale field of view.

Polarized light imaging (PLI) is a popular technique used for probing physiological structures in the brain with high spatial resolution, which has shown great promise for dMRI validation (Mehta et al. (2013)). In particular, the myelin sheath surrounding neurons is strongly birefringent (Morgan et al. (2021)), which will cause a phase delay in polarized light along the axis as the neuronal fibers. Traditionally, PLI is conducted by transmitting light of a specific polarization through a thin (on the order of 100 microns) slice of fixed tissue. The polarized light will interact with the cellular and sub-cellular structures, altering the polarization state, which is then measured by analyzing the transmitted light with a polarizer and camera detector. Rotating the analyzing polarizer in discrete increments and measuring the resulting transmitted power enables the estimation of the magnitude birefringence (also called “retardance”) and the orientation of the underlying structures (the “retardance angle”) (Larsen et al. (2007)). While valuable for mapping out fiber orientation, this approach is limited to only measuring specimens that are thin, fixed, and sectioned. Moreover, retardance and retardance angle are only two of many different polarization properties that could encode valuable information about the underlying tissue microstructure (Ghosh et al. (2008, 2009)).

Backscattering PLI, in particular those based on Mueller Matrix or Stokes polarimetry is emerging as a valuable technique for comprehensive polarization characterization of tissue specimens. Multiple approaches exist for Mueller Matrix polarimetry but in general, an optical system is constructed to cycle through different input polarization states and sample different output polarization states (Qi et al. (2018)). The collected data can then be used to reconstruct the Mueller Matrix, which provides a complete picture of the polarization properties of the material. The Mueller Matrix can be further decomposed into individual elements such as the linear retardance, diattenutation and circular polarization (Ghosh and Vitkin (2011)). Some recent works have shown that multi-wavelength Stokes polarimetry has great promise for brain microstructural mapping (Menzel et al. (2015); Borovkova et al. (2022)), but current studies remain limited to sectioned specimens. There remains a need to advance this technology toward *in vivo* acquisition, the first step of which is to demonstrate the utility of the technique on unsectioned whole brain specimens.

In this brief research report, we present a preliminary study to test the feasibility of multi-wavelength backscattering Mueller Matrix polarization microscopy for microstructural characterization of whole brain specimens in reflectance mode. For three specimens of ferret brain, we measure three different regions of the brain with known microstructural geometry: two composed of coherent white matter, and one composed of grey matter. We demonstrate that the technology is sensitive to orientation and fiber coherence in white matter, and shows promise for measuring grey matter microstructure. These results motivate further study of backscattering Mueller Matrix polarimetry in the context of *in vivo* characterization of brain microstructure, particularly for grey matter regions.

## 2 MATERIALS AND METHODS

### 2.1 Specimen Preparation

Perfusion-fixed ferret brain specimens (n=3) were generated in a previous unrelated study and provided for whole brain imaging in this study (Hutchinson et al. (2017)). Briefly, adult male ferrets underwent perfusion fixation with 4% paraformaldehyde (PFA). The brains were removed and post-fixed in 4% PFA for 10 days and then transferred for re-hydration and long-term storage under refrigeration in phosphate buffered saline with 0.01% sodium azide.

### 2.2 Polarization Imaging

For fixed brain specimens (Figure 1A), images were collected for three regions with known physiological and microstructural characteristics using a benchtop multispectral polarization microscope. Images were acquired of the corticospinal tract (CST), the optic chiasm (OC) and the cerebellum. These regions were selected due to their well known microstructural orientation. The corticospinal tract is a one-directional white matter fiber bundle along the direction of the spinal tract; the optic chiasm is composed of two, crossing white matter fiber bundles; and the cerebellum is a region of grey matter, with parallel fibers located beneath the surface layer. Thus, these three regions provide increasing complexity to examine the capabilities of the polarization imaging technology.

**Figure 1.**
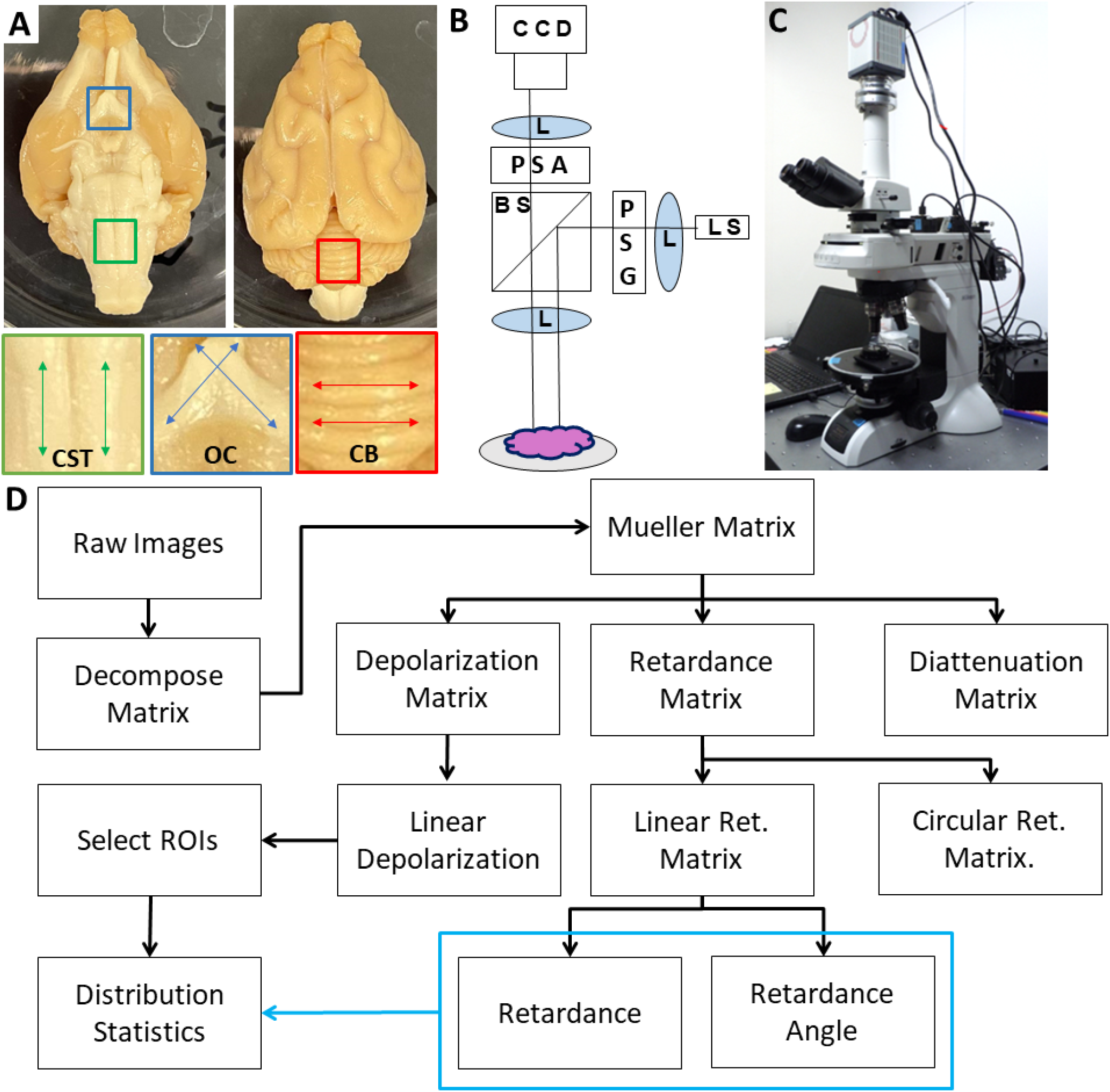
(A) Photograph of Ferret brain specimen along with regions of interest including corticospinal tract (CST); optic chiasm (OC) and cerebellum (CB). (B) Schematic diagram of the PLI system where light is emitted from a multi-wavelength light source (LS) which passes through a lens (L) and polarization state generator (PSG) before being directed to the sample through a beamsplitter (BS) and imaging lens. Backscattered light is collected and directed through a polarization state analyzer (PSA) before being focused onto a CCD camera. (C) Photograph of the polarimetry system. (D) Flow chart of data analysis including Mueller Matrix decomposition and calculation of distribution statistics.

The polarization microscope is a custom instrument developed by Nikon (Oka and Saito (2006); Saito et al. (2013)), which consists of an episcopic illumination system capable of dark-field illumination in five wavelengths and a polarimetric imaging system for measuring the state of polarization of scattered light from the tissue sample positioned at the microscope’s specimen stage (Figure 1B,C). A Nikon bright-field objective lens with minimal polarization distortion (Nikon XFI TU Plan Fluor EPI) was used to operate within the epi-illumination and dark-field imaging constraints of this polarimeter. 5X magnification was used to capture all images due to its larger field of view and depth of field. The instrument has been previously characterized, validated, and used for human tissue imaging, and produces spatial pixel-wise mappings of the sixteen Mueller Matrix components (Fujii et al. (2019); Saito et al. (2019)). Exposure times were optimized using the instruments “auto-adjust” setting.

### 2.3 Image Processing and Analysis

Wavelengths where the Mueller Matrix was unsuccessfully reconstructed were excluded from analysis. Incomplete reconstruction can be caused by tissue-dependent saturation, numerical instability, or other sources of noise. Ultimately, the 405 nm and 442 nm were analyzed for the CST, all wavelengths except 442 nm were analyzed for the OC and all five wavelengths were analyzed for the cerebellum. The raw Mueller Matrix maps were decomposed using the Lu-Chipman decomposition to extract polarization parameters including retardance, and retardance angle (Figure 1D) (Lu and Chipman (1996)).

In order to quantify specific tissues of interest, for each region that was imaged a sample-specific region of interest (ROI) was selected using ImageJ with the depolarization image as a reference (Rasband (2012)). Each region of interest was saved as a separate binary mask. For the CST, an ROI was selected for the spinal tract and the surrounding background region (known as the olivary complex, which lacks significant white matter directionality); for the OC, an ROI was selected for each of the two, crossing, coherent white matter bundles (referred to as left and right lobes), and for the cerebellum, a single ROI was selected for a region of the surface with minimal curvature.

Pixel-wise distributions of retardance and retardance angle were generated to qualitatively assess the polarizing characteristics of each ROI. For the CST, the total retardance was compared between the CST and background by taking the mean value of the distribution for each sample and averaging across the collected samples. We characterize the distributions of the retardance and retardance angle by measuring the mean, standard deviation, and kurtosis and taking the ratio of the CST to the background. For the OC, we compare the mean retardance of the distributions, as well as the difference in mean retardance angle between the different samples. For the cerebellum, we quantify the mean, standard deviation, and kurtosis of the retardance and retardance angle distributions as a function of wavelength.

### 2.4 Statistical Analysis

Statistical analysis of the quantitative comparisons for retardance and retardance angle for CST and OC was done using a paired t-test, as each sample had two comparable regions of interest. The cerebellum only had one region of interest and therefore no statistical comparisons were conducted. For the OC, we also extracted a radial slice and examined the retardance angle along this slice. This was done to examine the response of the imaging technique in a region of crossing fibers. To determine if the response is produced as an averaging of the microstructure or responding to the dominant orientation, we fit the resulting data along the slice to a linear curve. If backscattering PLI is sensitive to the averaging of the microstructure, the retardance angle would manifest as a linear gradient across the crossing fiber region; alternatively, sensitivity to the dominant microstructure would manifest as a step-function between the two crossing fiber directions. To test the linearity of this relationship, we conduct a hypothesis test with a null hypothesis that the slope is zero by using a Wald Test with t-distribution of the test statistic.

## 3 RESULTS

### 3.1 Corticospinal Tract

Figure 2 shows the results for measurements of the CST for whole brain specimens. Of the five wavelengths used for imaging, only 405 nm and 445 nm were unsaturated with the optimized exposure times. Figure 2A is a photograph of the specimen, while Figure 2B and 2C show the retardance and retardance angle maps for the region of interest showing the CST against the background olivary region (BKG). Quantification of these maps (Figure 2D,E) shows that there is a significant increase in retardance for the CST compared to the background (p<0.05) for both 405 nm and 442 nm. The retardance distributions for both tissue types resemble a normal distribution with similar spreads, but the CST has a significantly higher mean retardance. Distributions of retardance angle (Figure 2E) indicate that the CST has a significantly more coherent distribution of angles, which can be shown quantitatively by nearly a four-fold decrease of standard deviation (p<0.05 at 405 nm and 442 nm), without any significant change in mean or kurtosis.

**Figure 2.**
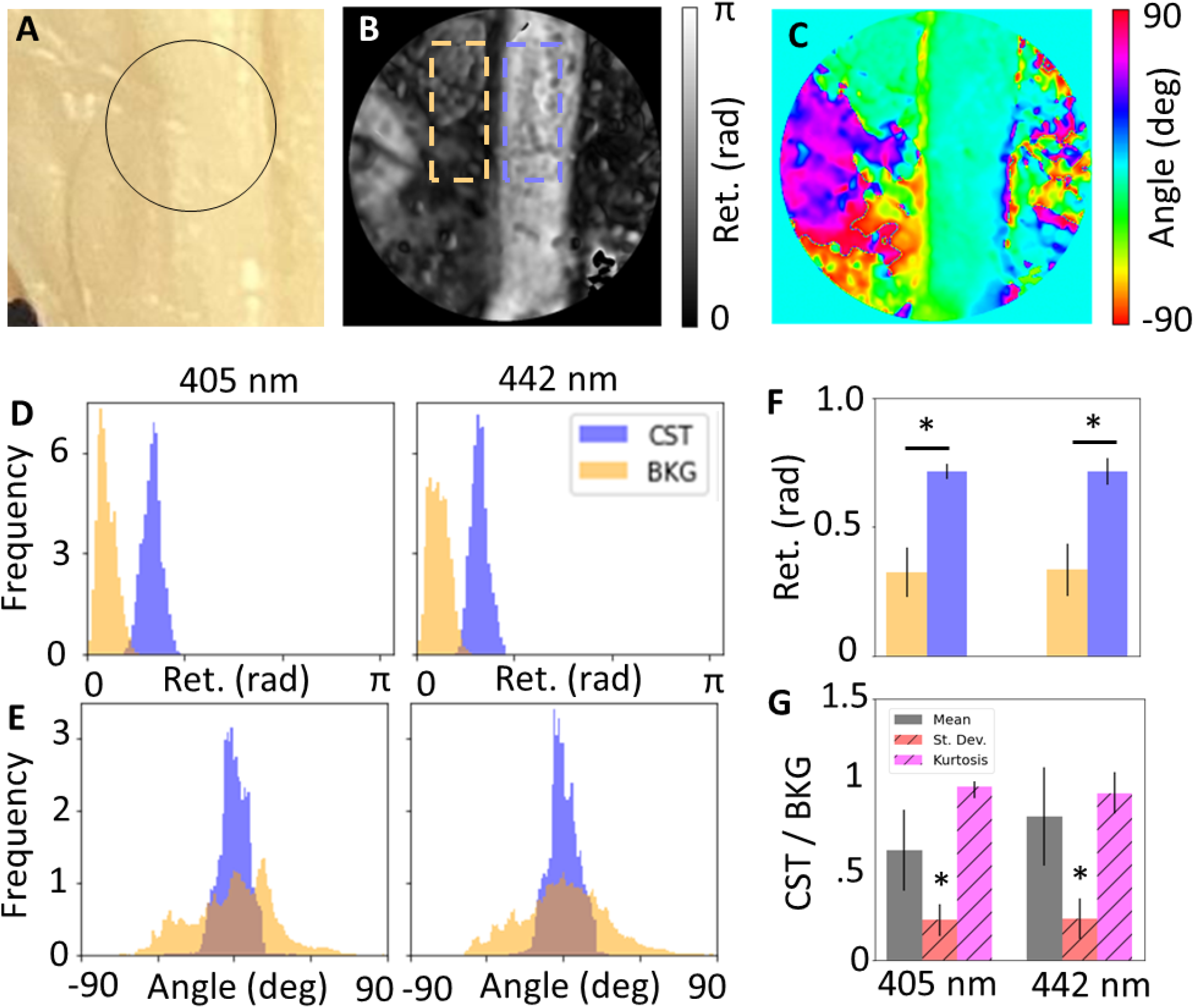
(A) Photograph of corticospinal tract (CST), maps of (B) retardance and (C) retardance angle of black circular region of interest shown in (A). Wavelength-resolved (D) retardance and (E) retardance angle distributions for CST compared to background (BKG). (F) Quantification of distribution statistics for retardance shows significance increase for CST compared to background, and (G) relative standard deviation and kurtosis of angle distributions shows significant decrease for CST, showing sensitivity to coherent fibers over unorganized tissue.

### 3.2 Optic Chiasm

Figure 3 shows the results for measurements of the OC for whole brain specimens. Only the 442 nm wavelength was excluded due to saturation and unsuccessful reconstruction of the Mueller Matrix. Figure 3A shows a photograph with the ROI outlines, while 3B and 3C show the retardance and retardance angle maps for the region of interest showing the left and right lobes. The retardance angle along the radial slice shown as a black curve in Figure 3C was extracted and the retardance angle was plotted against the radial angle (Figure 3D), which was found to have a linear slope (p<0.001), suggesting that the retardance angle is dictated by an average of the underlying microstructure, and not dictated by the dominant microstructural influence.

**Figure 3.**
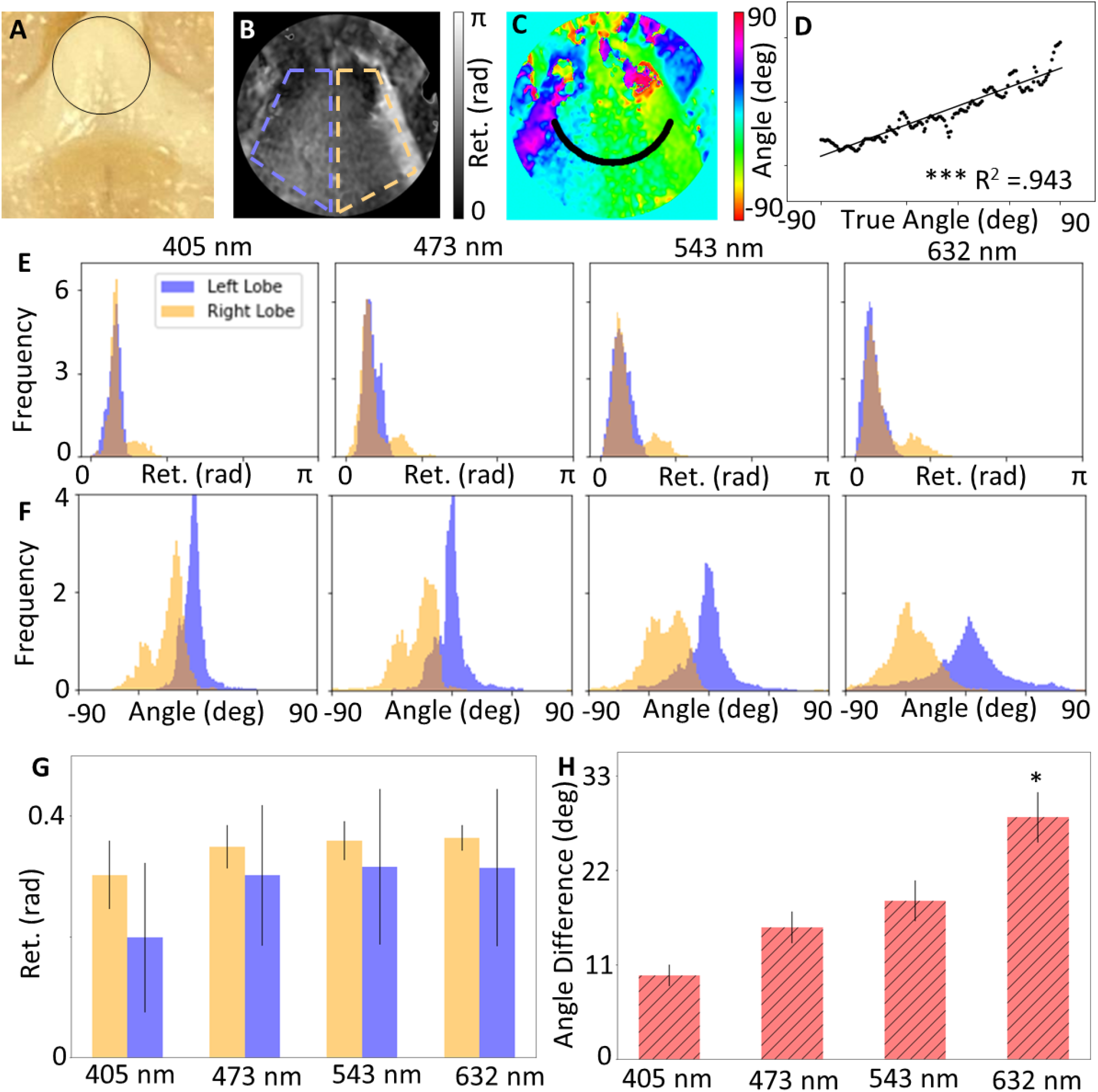
Photograph of optic chiasm (OC), maps of (B) retardance and (C) retardance angle of black circular region of interest shown in (A). (D) Circular slice of retardance angle along black line shown in (C), which shows a statistically significant linear relationship with radial angle. Wavelength-resolved (E) retardance and (F) retardance angle distributions for CST compared to background (BKG). (G) Quantification of mean retardance shows no significance difference for both lobes, and (H) the mean retardance angle for both lobes increases with wavelength.

Distributions of the retardance and retardance angle for the ROIs (Figure 3E, 3F) shows that the retardance of both lobes is approximately equal, which is expected given that both lobes are composed of white matter. However, the retardance angle distributions indicate that the difference in retardance angle becomes more prominent at higher wavelengths. This may be due to the higher penetration depth of longer wavelengths, enabling more sensitive measurements. Figures 3G and 3H show the quantification of these distributions, which support the qualitative observations; where there is no significant different in retardance across all wavelengths, but there is a statistically significant difference in the angle of the two lobes at a wavelength of 632 nm (p<0.05). Some distributions such as the right lobe in Figure 3E,F seem to exhibit bimoodal behavior, which may be caused by the influence of the curved geometry of the sample resulting in variations in fiber orientation.

### 3.3 Cerebellum

Figure 4 shows the results for the measurements of the cerebellum. Figure 4A is a photograph of the specimen, with Figure 4B demonstrating the expected underlying microstructural content of parallel fiber, which runs along the length of the ridges. Figures 4C and 4D show maps of the retardance and retardance angle, respectively. Distributions of the retardance and retardance angle for the ROI (Figure 4E, 4F) seems to suggest that the retardance is relatively constant across the tissue, but decreasing with higher wavelength. Conversely, the retardance angle seems to increase but become less tightly distributed. Quantification of the characteristics of these distributions through mean, standard deviation and kurtosis (Figure 4G, 4H) shows consistent observations - the mean retardance decreases with increasing wavelength and mean retardance angle increases with increasing wavelength. Given that the ridges are not oriented horizontally, we would expect a nonzero retardance angle if the measurements are sensitive to the underlying parallel fibers. These results seem to suggest that higher wavelengths may be more sensitive to the underlying microstructure, which could be due to the higher penetration depth. A further consideration regarding penetration depth is that an additional orientation feature may arise due to the climbing fibers that reach beneath the molecular layer of the cerebellum (Figure 4B).

**Figure 4.**
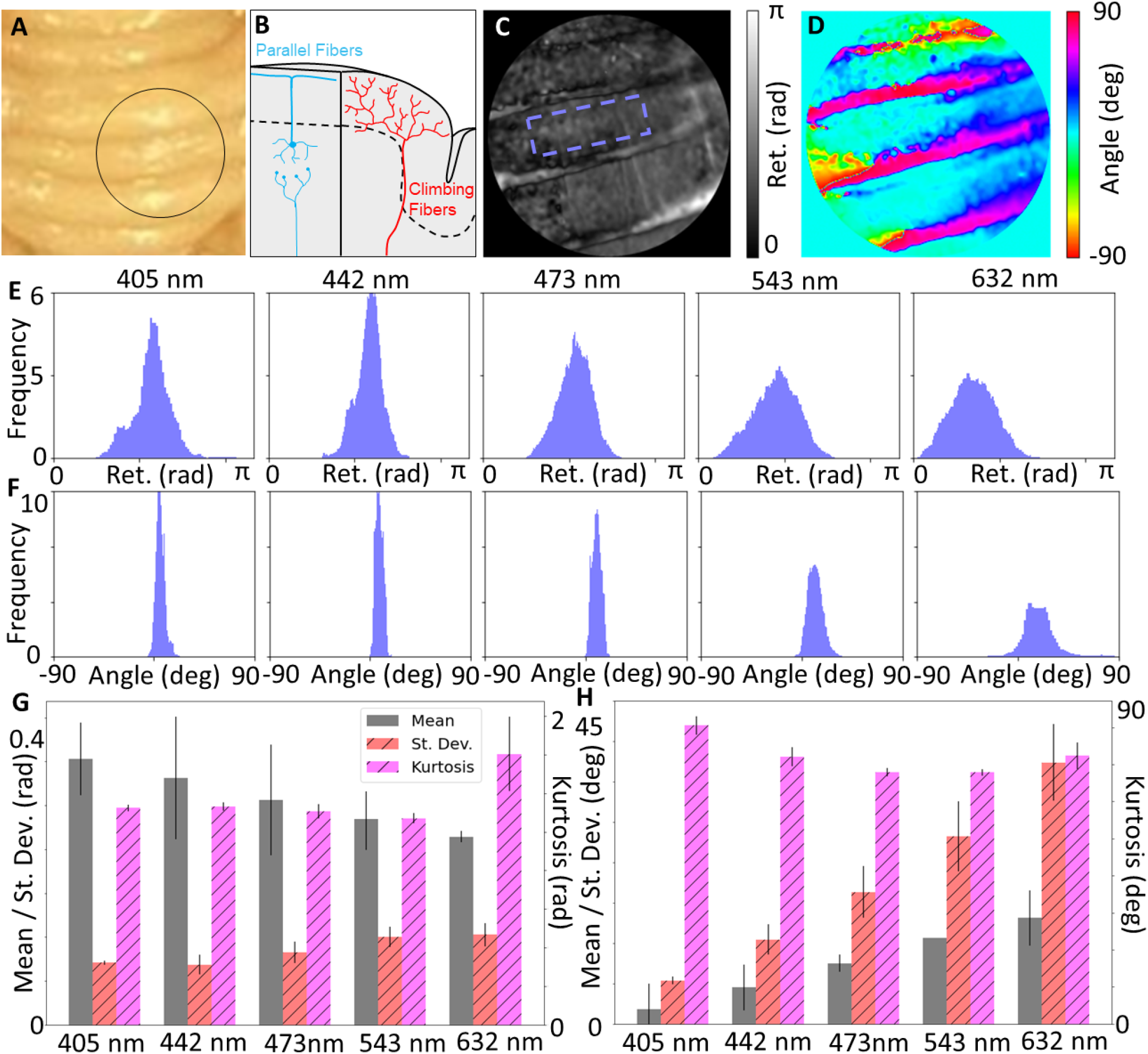
(A) Photograph of cerebellum, along with (B) diagram showing expected physiological structure of the grey matter. Maps of (C) retardance and (D) retardance angle of black circular region of interest shown in (A). Wavelength-resolved (E) retardance and (F) retardance angle distributions for ROI with minimal curvature. (G) Quantification of distributions statistics for retardance shows a decrease of the mean with increasing wavelength, and (H) retardance angle shows an increase of the mean with increasing wavelength, suggesting increased depth penetration of longer wavelengths may be more sensitive to parallel fiber structures.

## 4 DISCUSSION

Our preliminary results suggest that backscattering Mueller Matrix polarimetry could be a promising technique for noninvasive mapping of microstructural orientation features in the brain. We show that we are able to obtain measurements of fiber coherence and density (retardance), as well as fiber orientation (retardance angle) that are consistent with known microstructural characteristics. One advantage to the Mueller Matrix approach is that the measurement also provides information about other polarization parameters including depolarization, diattenuation, and circular polarization interaction. These properties may also contain valuable information for probing tissue microstructure, which could complement the known markers of retardance and retardance angle.

Primarily, other works investigating Mueller Matrix polarimetry in the context of brain microstructural imaging are limited to sectioned, fixed specimens of brain tissues (Borovkova et al. (2022); Schucht et al. (2020)). Our demonstration of this technology is valuable in that it evaluates the potential for *in vivo* use through the application to intact whole brain specimens in reflectance mode. However, other steps are necessary to conduct *in vivo* studies, for example optimizing acquisition times to mitigate motion artifacts. In addition, our study shows preliminary promise for using PLI to measure to grey matter microstructure, for which developing validation techniques is becoming increasing important as dMRI methods continue to advance. Further study is needed to validate the capabilities of this technology for grey matter microstructural assessment and likely hardware and image analysis algorithm optimizations are required to maximize the performance of the technology.

While promising, our results indicate that the selection of wavelength is critical. Wavelength-resolved light-tissue interaction is predominantly dictated by absorption, scattering, and penetration depth (Tuchin (2003)). Primary tissue absorbers are melanin, lipids and water, which give rise to an optical window in the visible spectrum (Smith et al. (2009)). Scattering will be influenced by wavelength and particle size but generally is higher for lower wavelengths, leading to decreased penetration depth. Given the heterogeneity of brain tissues, it may be necessary to optimize acquisition wavelength for specific anatomical structures and regions; we see that for the white matter regions such as the corticospinal tract, there is severe oversaturation at higher wavelength compared to the grey matter regions in the cerebellum. Our measurements may suggest that there is some sensitivity of polarized light to grey matter microstructure, as we observed an increased signature of retardance angle from the cerebellum, which is consistent with the angle of parallel fibers. This becomes more pronounced at higher wavelengths, which may be due to the requisite deeper penetration needed for the light to interact with the parallel fiber structures. In addition, the wavelength of light may influence the size of the tissue structures it primarily interacts with, providing more motivation to further understand how wavelength selection can be used as a tool to optimize the acquisition of microstructural information.

As anticipated, measuring non-flat geometries, such as with whole brain specimens, is particularly challenging and may introduce artifacts into the results. To account for this effect, further studies utilizing tissue phantoms will be undertaken to characterize how curved surfaces may influence the acquisition and reconstruction of Mueller Matrix data. The influence of topology may be correctable through calibration and data processing techniques; alternatively, hardware optimization could also be undertaken to increase the depth of field to enable in-focus imaging of samples with significant topological features. With a two-dimensional acquisition, the technology is only sensitive to in-plane orientation; further innovations could be made with the integration of three-dimensional acquisition through sample or hardware tilting, among other approaches Menzel et al. (2015); Borovkova et al. (2022).

## 5 CONCLUSION

Evaluation of microscale physiology and microstructural cellular features of the brain is becoming increasingly important to understand the mechanisms of neurodegenerative diseases, development and aging. While imaging modalities such as diffusion magnetic resonance imaging (dMRI) are extremely powerful for noninvasive, three-dimensional, microstructural characterization of tissues, there is a need for robust validation techniques to verify the biological accuracy of dMRI imaging data, particularly as these techniques are extended to probe grey matter. Polarization imaging is a promising technique for high resolution direct microstructural imaging, but application is typically limited to thin sectioned samples in transmission mode. We have shown for the first time that backscattering Mueller Matrix polarimetry in reflectance mode can be used to map microstructural features of intact brain tissue specimens with high biological accuracy. The experimental results show that backscattering polarimetry can probe white matter fiber coherence and fiber orientation in whole brain specimens, and show promise for probing grey matter microstructure. Ultimately, these preliminary results motivate further study to fully understand how backscattering polarimetry can best be used for validation of *in vivo* microstructural imaging of the brain.

## AUTHOR CONTRIBUTIONS

KM and RC contributed to data collection. JB assisted with data collection and performed image analysis and processing. CC, RC, and EH contributed to specimen preparation. TS and EH contributed to the study design. TS wrote the first draft of the manuscript. ND contributed to data collection and manuscript preparation. All authors contributed to manuscript revision, read and approved the submitted version.

## ACKNOWLEDGMENTS

We would like to thank the Nikon Corporation and Nikon Research Corporation of America for the use of the polarimeter system, as well as Saito Naooki and Heather Durko for technical guidance. We also would like to thank Faith Rice and Dr. Jennifer Barton for assistance with operating the polarimeter.

## REFERENCES

Bhagat, Y. A. and Beaulieu, C. (2004). Diffusion anisotropy in subcortical white matter and cortical gray matter: changes with aging and the role of CSF-suppression. Journal of magnetic resonance imaging: JMRI 20, 216–227. doi:10.1002/jmri.20102

Borovkova, M., Sieryi, O., Lopushenko, I., Kartashkina, N., Pahnke, J., Bykov, A., et al. (2022). Screening of Alzheimer’s Disease With Multiwavelength Stokes Polarimetry in a Mouse Model. IEEE transactions on medical imaging 41, 977–982. doi:10.1109/TMI.2021.3129700

Forouzannezhad, P., Abbaspour, A., Fang, C., Cabrerizo, M., Loewenstein, D., Duara, R., et al. (2019). A survey on applications and analysis methods of functional magnetic resonance imaging for Alzheimer’s disease. Journal of neuroscience methods 317, 121–140. doi:10.1016/j.jneumeth.2018.12.012

Fujii, T., Yamasaki, Y., Saito, N., Sawada, M., Narita, R., Saito, T., et al. (2019). Polarization characteristics of dark-field microscopic polarimetric images of human colon tissue. In Proc.SPIE. vol. 10890. doi:10.1117/12.2509000

Ghosh, N. and Vitkin, I. A. (2011). Tissue polarimetry: concepts, challenges, applications, and outlook. Journal of Biomedical Optics 16, 110801. doi:10.1117/1.3652896

Ghosh, N., Wood, M. F. G., Li, S.-h., Weisel, R. D., Wilson, B. C., Li, R.-K., et al. (2009). Mueller matrix decomposition for polarized light assessment of biological tissues. Journal of Biophotonics 2, 145–56. doi:10.1002/jbio.200810040

Ghosh, N., Wood, M. F. G., and Vitkin, I. A. (2008). Mueller matrix decomposition for extraction of individual polarization parameters from complex turbid media exhibiting multiple scattering, optical activity, and linear birefringence. Journal of Biomedical Optics 13, 044036. doi:10.1117/1.2960934

Guerrero-Gonzalez, J. M., Yeske, B., Kirk, G. R., Bell, M. J., Ferrazzano, P. A., and Alexander, A. L. (2022). Mahalanobis distance tractometry (MaD-Tract) - a framework for personalized white matter anomaly detection applied to TBI. NeuroImage 260, 119475. doi:10.1016/j.neuroimage.2022.119475

Gunning-Dixon, F. M., Brickman, A. M., Cheng, J. C., and Alexopoulos, G. S. (2009). Aging of cerebral white matter: a review of MRI findings. International journal of geriatric psychiatry 24, 109–117. doi:10.1002/gps.2087

Hutchinson, E. B., Schwerin, S. C., Avram, A. V., Juliano, S. L., and Pierpaoli, C. (2018). Diffusion MRI and the detection of alterations following traumatic brain injury. Journal of neuroscience research 96, 612–625. doi:10.1002/jnr.24065

Hutchinson, E. B., Schwerin, S. C., Radomski, K. L., Sadeghi, N., Jenkins, J., Komlosh, M. E., et al. (2017). Population based MRI and DTI templates of the adult ferret brain and tools for voxelwise analysis. NeuroImage 152, 575–589. doi:10.1016/j.neuroimage.2017.03.009

Lamptey, R. N. L., Chaulagain, B., Trivedi, R., Gothwal, A., Layek, B., and Singh, J. (2022). A Review of the Common Neurodegenerative Disorders: Current Therapeutic Approaches and the Potential Role of Nanotherapeutics. International journal of molecular sciences 23. doi:10.3390/ijms23031851

Larsen, L., Griffin, L. D., Grässel, D., Witte, O. W., and Axer, H. (2007). Polarized light imaging of white matter architecture. Microscopy research and technique 70, 851–863. doi:10.1002/jemt.20488

Le Bihan, D. (2014). Diffusion MRI: what water tells us about the brain. EMBO molecular medicine 6, 569–573. doi:10.1002/emmm.201404055

Liu, Y., Lu, L., Li, F., and Chen, Y.-C. (2022). Neuropathological Mechanisms of Mild Traumatic Brain Injury: A Perspective From Multimodal Magnetic Resonance Imaging. Frontiers in neuroscience 16, 923662. doi:10.3389/fnins.2022.923662

Lu, S.-Y. and Chipman, R. A. (1996). Interpretation of Mueller matrices based on polar decomposition. Journal of the Optical Society of America A 13, 1106. doi:10.1364/JOSAA.13.001106

Madden, D. J., Bennett, I. J., and Song, A. W. (2009). Cerebral white matter integrity and cognitive aging: contributions from diffusion tensor imaging. Neuropsychology review 19, 415–435. doi:10.1007/s11065-009-9113-2

Mehta, S. B., Shribak, M., and Oldenbourg, R. (2013). Polarized light imaging of birefringence and diattenuation at high resolution and high sensitivity. Journal of optics (2010) 15. doi:10.1088/2040-8978/15/9/094007

Menzel, M., Michielsen, K., De Raedt, H., Reckfort, J., Amunts, K., and Axer, M. (2015). A Jones matrix formalism for simulating three-dimensional polarized light imaging of brain tissue. Journal of the Royal Society, Interface 12, 20150734. doi:10.1098/rsif.2015.0734

Minchew, H. M., Ferren, S. L., Christian, S. K., Hu, J., Keselman, P., Brooks, W. M., et al. (2022). Comparing imaging biomarkers of cerebral edema after TBI in young adult male and female rats. Brain research 1789, 147945. doi:10.1016/j.brainres.2022.147945

Morgan, M. L., Brideau, C., Teo, W., Caprariello, A. V., and Stys, P. K. (2021). Label-free assessment of myelin status using birefringence microscopy. Journal of neuroscience methods 360, 109226. doi:10.1016/j.jneumeth.2021.109226

Nguyen, A. T., Kouri, N., Labuzan, S. A., Przybelski, S. A., Lesnick, T. G., Raghavan, S., et al. (2022). Neuropathologic scales of cerebrovascular disease associated with diffusion changes on MRI. Acta neuropathologica doi:10.1007/s00401-022-02465-w

Novikov, D. S., Fieremans, E., Jespersen, S. N., and Kiselev, V. G. (2019). Quantifying brain microstructure with diffusion MRI: Theory and parameter estimation. NMR in biomedicine 32, e3998. doi:10.1002/nbm.3998

Oka, K. and Saito, N. (2006). Snapshot complete imaging polarimeter using Savart plates. In Proc.SPIE. vol. 6295. doi:10.1117/12.683284

Pfefferbaum, A., Adalsteinsson, E., Rohlfing, T., and Sullivan, E. V. (2010). Diffusion tensor imaging of deep gray matter brain structures: effects of age and iron concentration. Neurobiology of aging 31, 482–493. doi:10.1016/j.neurobiolaging.2008.04.013

Qi, J., He, H., Lin, J., Dong, Y., Chen, D., Ma, H., et al. (2018). Assessment of tissue polarimetric properties using Stokes polarimetric imaging with circularly polarized illumination. Journal of biophotonics 11, e201700139. doi:10.1002/jbio.201700139

Rasband, W. S. (2012). ImageJ. U. S. National Institutes of Health, Bethesda, Maryland, USA, //imagej.nih.gov/ij/

Rathi, Y., Pasternak, O., Savadjiev, P., Michailovich, O., Bouix, S., Kubicki, M., et al. (2014). Gray matter alterations in early aging: a diffusion magnetic resonance imaging study. Human brain mapping 35, 3841–3856. doi:10.1002/hbm.22441

Rosenkranz, M. A., Dean, D. C. r., Bendlin, B. B., Jarjour, N. N., Esnault, S., Zetterberg, H., et al. (2022). Neuroimaging and biomarker evidence of neurodegeneration in asthma. The Journal of allergy and clinical immunology 149, 589–598. doi:10.1016/j.jaci.2021.09.010

Saito, N., Odate, S., Otaki, K., Kubota, M., Kitahara, R., and Oka, K. (2013). Wide field snapshot imaging polarimeter using modified Savart plates. In Proc.SPIE. vol. 8873. doi:10.1117/12.2022829

Saito, N., Sato, K., Fujii, T., Durko, H. L., Goldstein, G. L., Phillips, A. H., et al. (2019). Multispectral Mueller matrix imaging dark-field microscope for biological sample observation. In Proc.SPIE. vol. 10890. doi:10.1117/12.2508109

Schucht, P., Lee, H. R., Mezouar, H. M., Hewer, E., Raabe, A., Murek, M., et al. (2020). Visualization of White Matter Fiber Tracts of Brain Tissue Sections With Wide-Field Imaging Mueller Polarimetry. IEEE transactions on medical imaging 39, 4376–4382. doi:10.1109/TMI.2020.3018439

Smith, A. M., Mancini, M. C., and Nie, S. (2009). Bioimaging: second window for in vivo imaging. Nature nanotechnology 4, 710–711. doi:10.1038/nnano.2009.326

Tuchin, V. V. (2003). Light-tissue interactions. In Biomedical Photonics Handbook (Boca Raton, Florida: CRC Press). 1–27

Wang, Z., Xin, J., Wang, Z., Yao, Y., Zhao, Y., and Qian, W. (2021). Brain functional network modeling and analysis based on fMRI: a systematic review. Cognitive neurodynamics 15, 389–403. doi:10.1007/s11571-020-09630-5

Zhao, H., Tsai, C.-C., Zhou, M., Liu, Y., Chen, Y.-L., Huang, F., et al. (2022). Deep learning based diagnosis of Parkinson’s Disease using diffusion magnetic resonance imaging. Brain imaging and behavior 16, 1749–1760. doi:10.1007/s11682-022-00631-y

